# Unexpected suppression of neural responses to natural foreground versus background sounds in auditory cortex

**DOI:** 10.1101/2023.11.20.567922

**Authors:** Gregory R. Hamersky, Luke A. Shaheen, Mateo López Espejo, Jereme C. Wingert, Stephen V. David

## Abstract

In everyday hearing, listeners encounter complex auditory scenes containing overlapping sounds that must be grouped into meaningful sources, or streamed, to be perceived accurately. A common example of this problem is the perception of a behaviorally relevant foreground stimulus (speech, vocalizations) in complex background noise (environmental, machine noise). Studies using a foreground/background contrast have shown that high-order areas of auditory cortex in humans pre-attentively form an enhanced representation of the foreground over background stimulus. Achieving this invariant foreground representation requires identifying and grouping the features that comprise the background noise so that they can be removed from the representation of the foreground. To study the cortical computations underlying representation of concurrent background (BG) and foreground (FG) stimuli, we recorded single unit responses in the auditory cortex (AC) of ferrets during presentation of natural sound excerpts from these two categories. In primary and secondary AC, we found overall suppression of responses when BGs and FGs were presented concurrently relative to the sum of responses to the same stimuli in isolation. Surprisingly, and in contrast to percepts that emphasize dynamic FGs, responses to FG sounds were suppressed relative to the paired BG sound. The degree of relative FG suppression could be explained by spectro-temporal statistics unique to each natural sound. Moreover, systematic degradation of the same spectro-temporal features decreased FG suppression as the sound categories became progressively less statistically distinct. The strongly suppressed representation of FG sounds in single units of AC in the presence of BG sound reveals a novel insight into how complex acoustic scenes are encoded at early stages of auditory processing.

## Introduction

When interacting with the world, listeners encounter auditory scenes containing dynamic, spectrally overlapping sounds. Accurate hearing in noisy environments requires streaming, that is, the grouping of sound features into representations of their distinct sources^1,2^. Because natural sounds are spectrally and temporally dynamic, they cannot be differentiated based on tonotopic channels inherited from the cochlea. Instead, streaming must distinguish stimuli according to statistical regularities in the time and frequency domains^1,3–7^. Streaming is engaged by active, top-down processes, such as during selective attention to the voice of a single speaker^8–11^, and pre-attentively, as during perception of behaviorally relevant stimuli in noise^12–15^. In both cases, the representation of one stimulus is selectively enhanced relative to others, both perceptually and in cortical activity. The computational processes by which neural responses to competing stimuli at early stages of auditory processing are identified and suppressed to produce this invariant representation remain poorly understood.

Psychoacoustic experiments have classically studied auditory streaming by identifying perceptually grouped sound features^16–29^. Such approaches often employ synthetic stimuli with parametric properties to precisely manipulate and probe the boundaries of streaming. Results from these studies highlight the importance of spectral^18,19,27,28^, temporal^20–22,29^, and spatial^19,23,30^ sound statistics to successful streaming. Natural stimuli are more complex and dynamic, but similar principles can be applied to model the streaming of natural sounds^13,31–33^.

One useful framework to study the streaming of natural sounds is to use a background/foreground (BG/FG) contrast. Many behaviorally relevant stimuli (e.g., speech and other vocalizations) are perceived preferentially over noisy backgrounds (wind, water, machinery, etc.)^14,15,34–36^. Consistent with this behavioral observation, local field potential (LFP) and functional imaging (fMRI) studies of the human superior temporal gyrus have shown that activity evoked by foreground sounds is largely invariant to background noise, suggesting that the auditory cortex intrinsically supports streaming of foregrounds over backgrounds^14,15^. While the distinction between background and foreground can be subjective and context-dependent, most sounds can be classified as one or the other based on quantitative properties of the sound spectrogram^4,34–37^.

Studies using single unit recordings in animal models have argued for noise-robust representations in primary auditory cortex (A1) as well^13,38,39^. This work has focused largely on simple, static noise rather than natural backgrounds. Moreover, as with much prior work in human STG^8,13–15^, most single-unit studies have used stimulus reconstruction methods. These approaches demonstrate the ability to recover the foreground stimulus but do not exclude the possibility that information is also preserved about background noise^40–42^. A small number of studies measuring evoked activity have indicated that information about both backgrounds and foregrounds can be represented at this earlier processing stage^43–45^. Thus, it remains uncertain if foreground representations are enhanced in A1 to a similar degree as in human STG, or if representations in A1 describe an intermediate representation of sound streams, supporting a noise-invariant foreground representation at later stages.

In the current study, we recorded single unit activity in the auditory cortex of awake, passively listening ferrets to understand how concurrent natural background and foreground sounds are represented. We used an experimental design in which natural background textures were presented concurrently with statistically distinct and more dynamic foreground sounds, serving as natural signals in noise. We report an unexpectedly dominant suppression of the foreground relative to background in the neural response, challenging intuitions about the ubiquitousness of robust noise-invariant representation of behaviorally salient stimuli across the auditory hierarchy.

## Results

### Unexpected suppression of cortical responses to natural foreground sounds by natural backgrounds

To investigate how neurons in auditory cortex (AC) integrate information about concurrent natural sounds, we recorded single-unit activity from passively listening, head-fixed ferrets (Figure 1A). Natural sound stimuli were drawn from two broad, ethologically relevant categories: background textures (BGs) and foreground transients (FGs). We used this BG/FG contrast based on previous work showing that this stimulus configuration produces streaming, with enhanced perception and cortical representation of the FGs over noisy BGs^13–15,38,39^.

**Figure 1.**
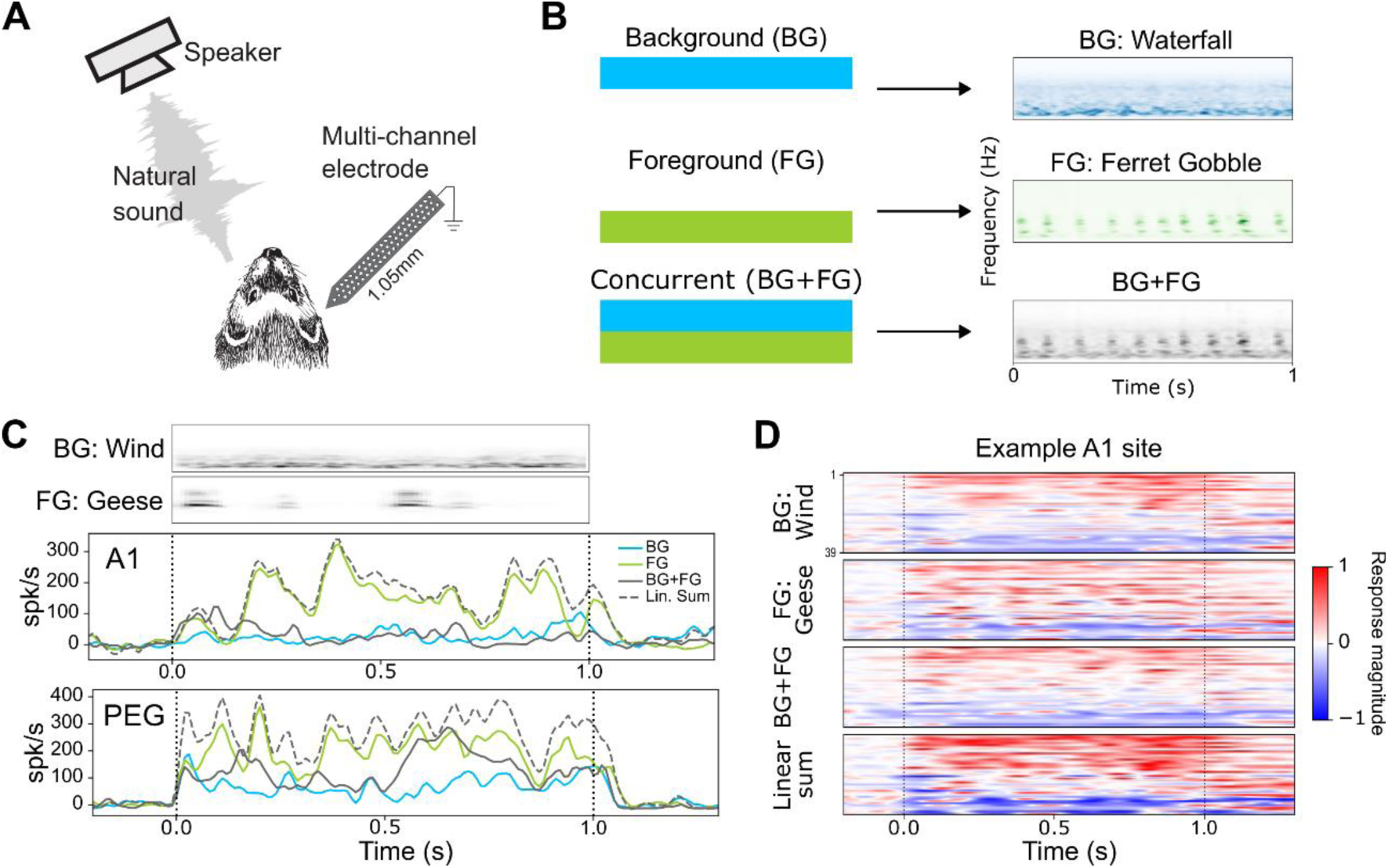
Characterization of neural responses to concurrent natural background (BG) and foreground (FG) sounds in ferret auditory cortex (AC) (A) Head-fixed ferrets were presented natural sound stimuli from a free-field speaker 30° contralateral to the recording hemisphere. Multi-channel microelectrode arrays recorded single-unit activity from primary (A1) or secondary (PEG) fields of AC. (B) During recording, 1s natural sound excerpts from two distinct, ethological categories—backgrounds (BGs) and foregrounds (FGs)—were presented in isolation (blue and green spectrograms, respectively) and concurrently (black spectrogram). (C) Example PSTH responses to the same BG/FG sound pairing from a unit in A1 (*upper*) and PEG (*lower*). In both cases, the linear sum (dashed) of the BG (blue) and FG (green) responses is greater than the actual BG+FG (black) response, indicating a suppressed BG+FG response. PSTHs were smoothed for visualization. (D) Heatmaps show normalized PSTH responses for all units (n = 39) in the A1 recording site shown in (C). The BG+FG panel has much lower response magnitude than the Linear sum, demonstrating the prevalence of suppressed, sub-linear BG+FG responses in most units.

At the beginning of each experiment, a set of 29 BGs and 41 FGs (1s duration) were presented in isolation, and the 3-5 sounds from each category that evoked the largest average multi-unit response were used in the subsequent recordings. Pairs of the selected BG and FG sounds were presented both individually and simultaneously (BG+FG, Figure 1B). We recorded 1,669 auditory-responsive units in A1 (2,766 units total) and 864 auditory-responsive units in periectosylvian gyrus (PEG), a secondary auditory field of auditory cortex (1,608 units total). Multiple FG/BG pairs were presented during each experiment. Individual units responded to at least one stimulus in subsets of the presented pairs, leading to a total of 14,360 responsive neuron/sound pairs in A1 (29,445 total) and 6,290 responsive neuron/sound pairs in PEG (14,936 total). Peristimulus time histogram (PSTH) responses to each stimulus were computed by averaging across 10-20 repetitions (Figure 1C).

To gain a basic understanding of how AC neurons respond to concurrent natural stimulus pairs, we first evaluated response linearity. We compared the evoked PSTH response to each concurrent BG+FG stimulus to the sum of responses to the BG and FG stimuli in isolation. Examples from both A1 and PEG show that evoked responses to BG+FG combinations were consistently lower than the sum of the responses to same sounds in isolation (Figure 1C). This suppression of BG+FG responses was observed across most recorded units (Figure 1D, Figure S1).

Given the observation of overall nonlinear suppression, our next question was how the component BG and FG stimuli contribute to the concurrent BG+FG response. Example responses to BG+FG stimuli were sometimes invariant to one of the stimuli (Figure 1C, *upper*), but in other cases they appeared to be a combination of the responses to the individual BG and FG stimuli (Figure 1C, *lower*). To quantify how each component contributed to the BG+FG response, we fit a model for the concurrent response as a linear weighted sum of the constituent responses:

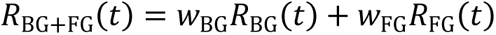

Weights were fit for each neuron and stimulus pair, minimizing the mean-squared error prediction of the actual BG+FG response (Figure 2A). Weights <1 indicate suppression relative to the component response. We compared weights between BG and FG categories for each neuron and stimulus pair tested. For A1 neuron/stimulus pairs with reliable responses to both individual BG and FG sounds (BG^+^/FG^+^, n = 9,057/29,445, see *Methods*, Figure S2A), weights were categorically divergent. Average *w_FG_* (median = 0.289 ± 0.002) was significantly lower than *w_BG_* (median = 0.663 ± 0.004, Wilcoxon signed rank test, p < 10^-9^), indicating a strong preferential suppression of FG responses (Figure 2B, 2C). Only neuron/stimulus pairs with a good model fit (r ≥ 0.4) were considered in analyses, but the same trend was observed for units with worse model performance (see *Methods*, Figure S2B). In PEG, neuron/stimulus pairs in the BG^+^/FG^+^ group (n = 3,838/14,936, Figure S2D), with r ≥ 0.4 (Figure S2E) showed a similarly strong preferential suppression of FG responses (*w_FG_:* median = 0.314 ± 0.005, *w_BG_:* median = 0.632 ± 0.004, Wilcoxon signed rank test, p < 10^-9^, Figure 2D).

**Figure 2.**
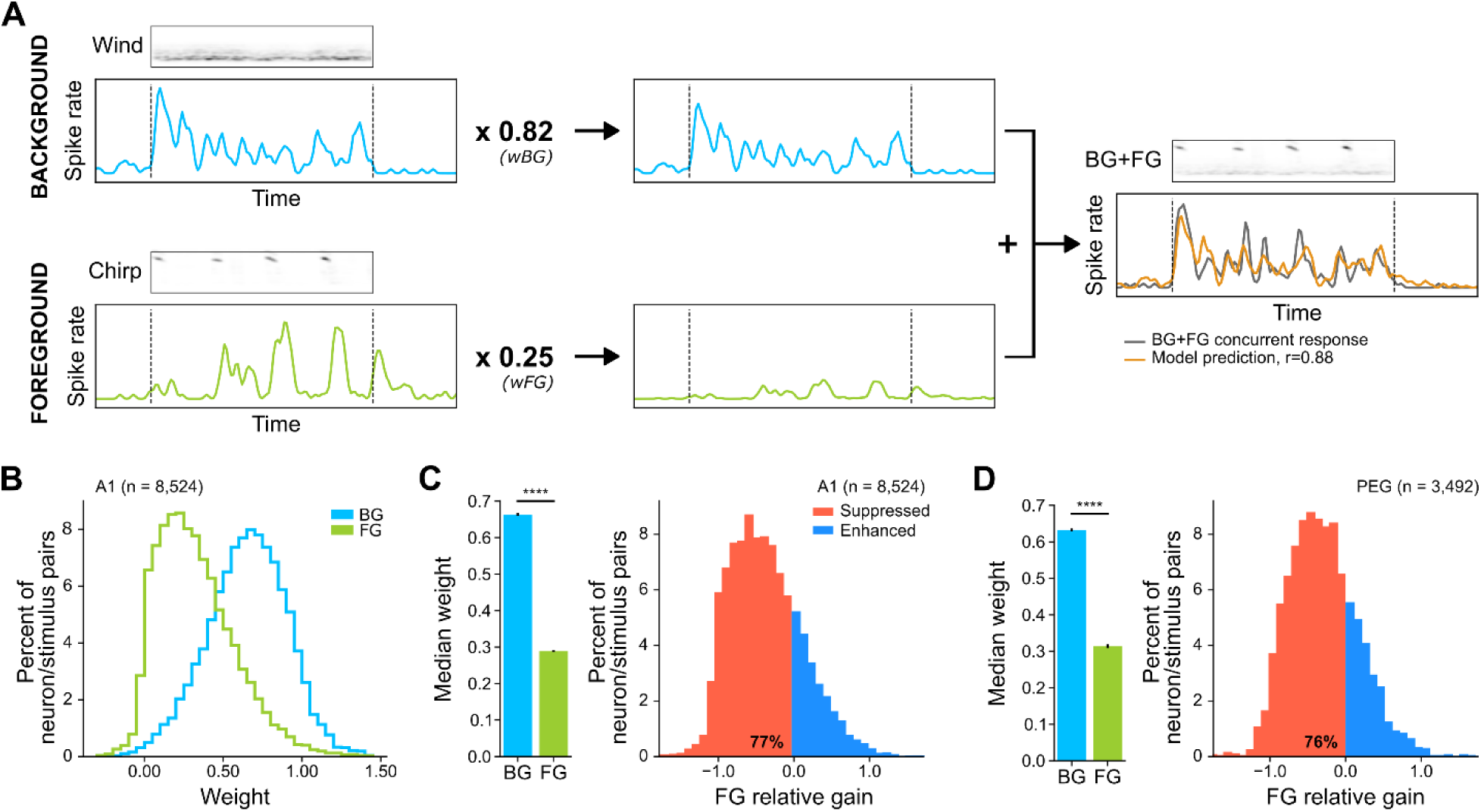
FG responses are preferentially suppressed relative to BG responses. (A) Diagram of the linear weighted model used to quantify the contribution of component BG and FG responses to concurrent BG+FG sound presentations. The PSTH response to each BG and FG sound in isolation was weighted and summed to minimize the mean-squared error prediction of the actual BG+FG response. (B) Histograms show distribution of BG (blue) and FG (green) weights for all neuron/sound pair combinations in A1. (C) Bars at left compare median weights (± jack-knifed S.E. across neuron/sound pairs, n = 8,532, Wilcoxon signed-rank test, ****p < 10^-9^). Histogram of FG relative gain (RG_FG_) for all A1 combinations. Negative RG_FG_ values (red) indicate neuron/sound pairs that show FG-specific suppression (77%). (D) (*left*) Median BG and FG weights in PEG (n = 3,501, ****p < 10^-9^) and (*right*) histogram of RG in PEG (RG_FG_ < 0 for 76%), plotted as in (C).

To verify that the weighted model recapitulated the general suppression reported above, we calculated the average weight, (*w_FG_ + w_BG_*) / 2, for each neuron/sound pair. The distribution of average weights significantly correlated with level of suppression observed in the response to the concurrent sounds, R*_BG+FG_* / (R*_BG_* + R*_FG_*) (A1: r = 0.67, p < 10^-9^, Figure S2C; PEG: r = 0.67, p < 10^-9^, Figure S2F). Thus, model weights provided a reasonable quantification of the overall and stimulus-selective suppression. This systematic FG suppression was not expected, given prior work indicating enhancement of responses to FG sounds in AC ^13–15,38,39^. Below, we describe several additional analyses to validate this result.

### Degree of FG suppression does not depend on selectivity for component stimuli

One possible explanation for FG suppression might be that neurons generally respond more strongly to BG stimuli and weigh the dominant stimulus in the concurrent BG+FG response. To determine whether FG suppression could be attributed to strong BG responses, we analyzed a subset of stimulus pairs where only one sound, BG or FG, evoked a response in isolation. We defined the BG^0^/FG^+^ and BG^+^/FG^0^ groups as neuron/stimulus pairs that responded only to the FG or BG stimulus, respectively (n = 3,117/29,445, 10.6%, n = 2,186/29,445, 7.4%, see *Methods,* Figure S2A). Comparison of *w_FG_* from the BG^0^/FG^+^ subset (median = 0.238 ± 0.006) and *w_BG_* from the BG^+^/FG^0^ subset (median = 0.605 ± 0.004) showed similar, significant (p < 10^-9^) preferential suppression of FGs in both A1 and PEG (*w_FG_:* median = 0.247 ± 0.010, *w_BG_:* median = 0.611 ± 0.011, p < 10^-9^, Figure S3B). The suppression of FG responses even in the absence of any BG-evoked response suggests that FG suppression can be driven by subthreshold activity. Thus, it does not simply reflect preferential responses for BG stimuli.

Given that sounds from both sound categories were suppressed relative to presentation in isolation (weight < 1), we combined the weights into a single metric, FG relative gain (RG_FG_), to describe the relative contribution of FG versus BG to the BG+FG response:

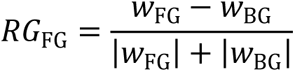

Values of RG_FG_ < 0 indicate suppression of FG relative to BG, which we refer to as FG-specific suppression. RG_FG_ values in A1 comprise a normal distribution centered significantly below zero (p < 10^-9^, Figure 2C, *right*). Consistent with weights observed for the BG^0^/FG^+^ and BG^+^/FG^0^ data, RG_FG_ depends only slightly on an individual neuron’s responsiveness to either the individual BG or FG stimulus (Figure S3A). Even when a neuron responded only to the FG in isolation, FG-specific suppression was usually observed. Thus, the suppression of FG responses appears to be categorical and dependent on activity in the wider AC network, rather than the tuning of a single neuron. This observation prompted further investigation into features of the BG versus FG stimuli that can explain this unexpected result.

### Neural responses adapt rapidly to concurrent stimulus presentations

Theories of natural sound streaming suggest that the brain computes statistical regularities over time in the neural population response to group sound features into perceptual objects^36,46^. To investigate the dynamics of BG+FG interactions, we measured the temporal window over which neural responses to a single sound adapt to the onset of a second sound. For a subset of recordings, we included stimulus instances where a BG played the entire 1s stimulus duration and the paired FG began 0.5s after BG onset (full BG + half FG; BG+hFG, Figure 3A) or vice versa (late BG onset, hBG+FG, Figure 3D). The late-onset stimuli were generated using the latter half of the full stimulus to allow direct comparison of responses to the second half of the standard BG+FG condition.

**Figure 3.**
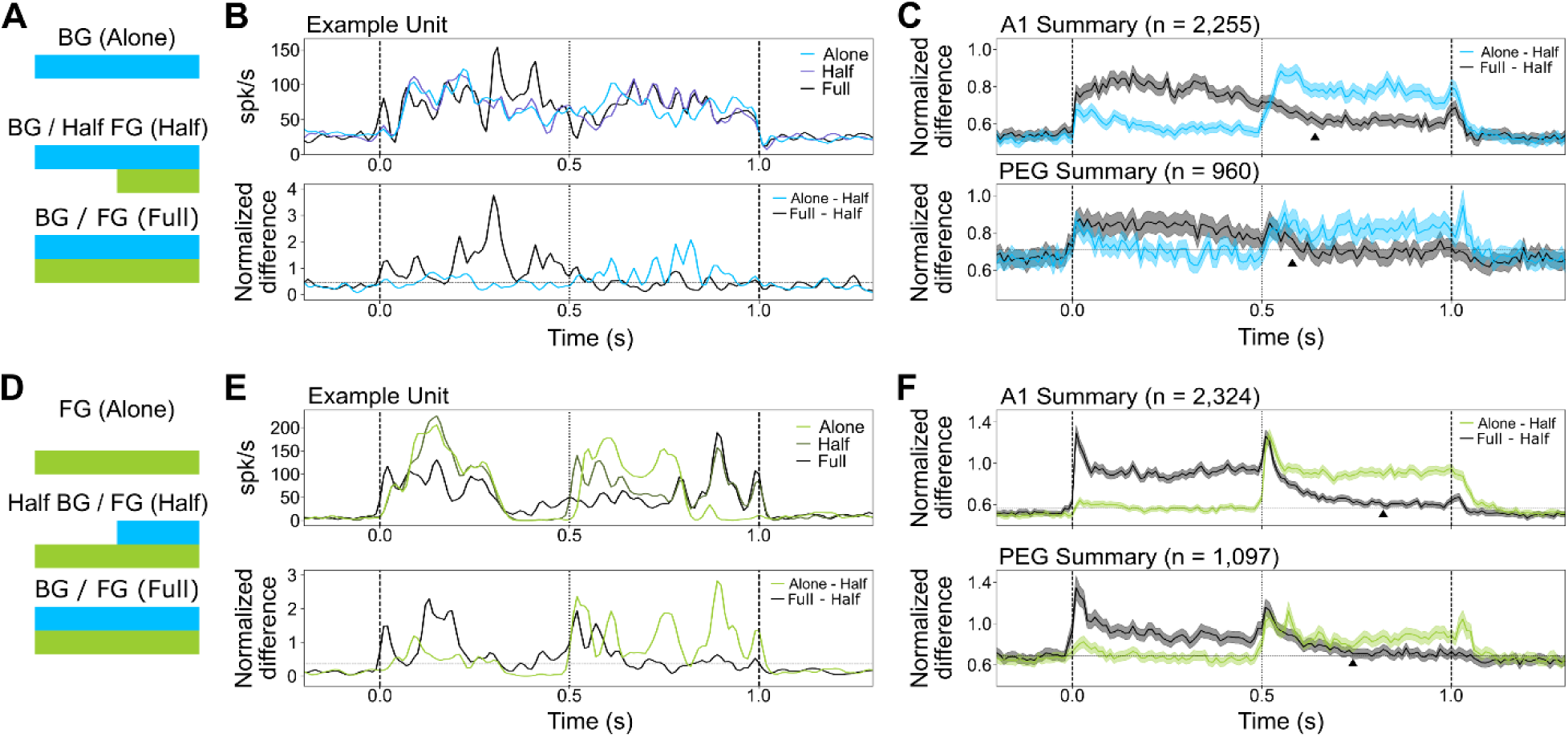
Response dynamics following concurrent sound onset differ between A1 and PEG. (A) Schematic of BG+hFG condition, in which a truncated 0.5s FG begins 0.5s after BG onset. (B) (*upper*) PSTH response of a single A1 neuron to BG+hFG, BG+FG, and BG alone stimuli. (*lower*) Difference in PSTH response between the BG+hFG condition and the BG+FG (black) and BG alone (blue). Values near zero indicate similar responses at the corresponding time point. (C) Normalized response difference between BG+hFG and the two reference conditions, averaged across all neuron/sound pair combinations in A1 (*upper*) and PEG (*lower*). Filled triangles indicate the latest time point at which the response following introduction of FG at 0.5s differs from the ongoing BG+FG response (A1: 140 ms, PEG: 80 ms). Significant differences were measured relative to a noise floor, computed by averaging BG+hFG and BG alone responses over 0.5 s. (D) Schematic of hBG+FG condition, where the BG sound is introduced 0.5 s following FG onset. (E) Example responses hBG+FG, BG+FG and FG alone stimuli, plotted as in (B). (F) Average normalized response difference for hBG+FG stimuli, plotted as in (C) (A1: 320 ms, PEG: 240 ms).

To measure response dynamics to the onset of an interrupting FG, we compared the half-FG (BG+hFG) response both to the full concurrent (BG+FG) response and to the BG alone response. Similarity of the BG+hFG response to the other conditions was computed as the difference in PSTH response, normalized by the standard deviation of each neurons’ time-varying spike rate across all stimuli. The first 0.5s of the BG+hFG and BG alone conditions are identical; thus, their difference should be minimal and provide a baseline error. After the half-FG onset at 0.5s, we expect the difference between the BG+hFG and BG+FG responses to decrease and converge to the baseline error. Example responses (Figure 3B) demonstrate the timing of this transition in PSTH responses, which indicates how long the response takes to adapt to the appearance of the FG sound. The converse analysis was performed by comparing the response to an interrupting BG (hBG+FG) to both the BG+FG and FG alone responses (Figure 3E).

Averaging the differential responses across all units and sound pairs for each condition allowed us to compute the average adaptation time following the onset of a second, concurrent stimulus. We defined the average adaptation time as the last time bin in which the response difference was significantly greater than the baseline error for three consecutive time bins (p < 0.05, jackknifed t-test). In the BG+hFG condition, adaptation in A1 took place more slowly (140 ms, n = 2,255) than in PEG (80 ms, n = 960, Figure 3C). Meanwhile, in the hBG+FG condition, which introduced the BG at 0.5 sec, adaptation times were overall longer, but the same trend was observed between A1 (320 ms, n = 2,324) and PEG (240 ms, n = 1,097, Figure 3F). Thus, adaptation was slower following the introduction of BG sounds, but, as in the case of FG sounds, it was faster in PEG than in A1.

### Degree of FG suppression is reduced over time

Example neurons suggest FG responses in A1 are suppressed throughout the duration of the BG+FG stimulus (Figure 1C, *upper*) while in PEG the degree of FG suppression can decrease over the course of the 1s stimulus (Figure 1C, *lower*). Based on our analysis of response dynamics (Figure 3), we determined that BG+FG responses should reach a steady state by 0.5s. To compare relative FG suppression before and after reaching a steady state, we repeated the relative weight analysis (Figure 2A) separately for the first and second 0.5s halves of the BG+FG response.

A1 and PEG both showed a significant decrease in FG suppression in the second half of the stimulus (A1: Wilcoxon signed rank test, p = 6.36e-8; PEG: p = 4.57e-7, Figure 4). RG_FG_ was not significantly different between A1 and PEG in the 0-0.5s fit period (p = 0.126). However, consistent with results showing more rapid adaptation dynamics in PEG, FG suppression in PEG during the 0.5-1s fit period was significantly decreased compared to A1 (p = 3.94e-3).

**Figure 4.**
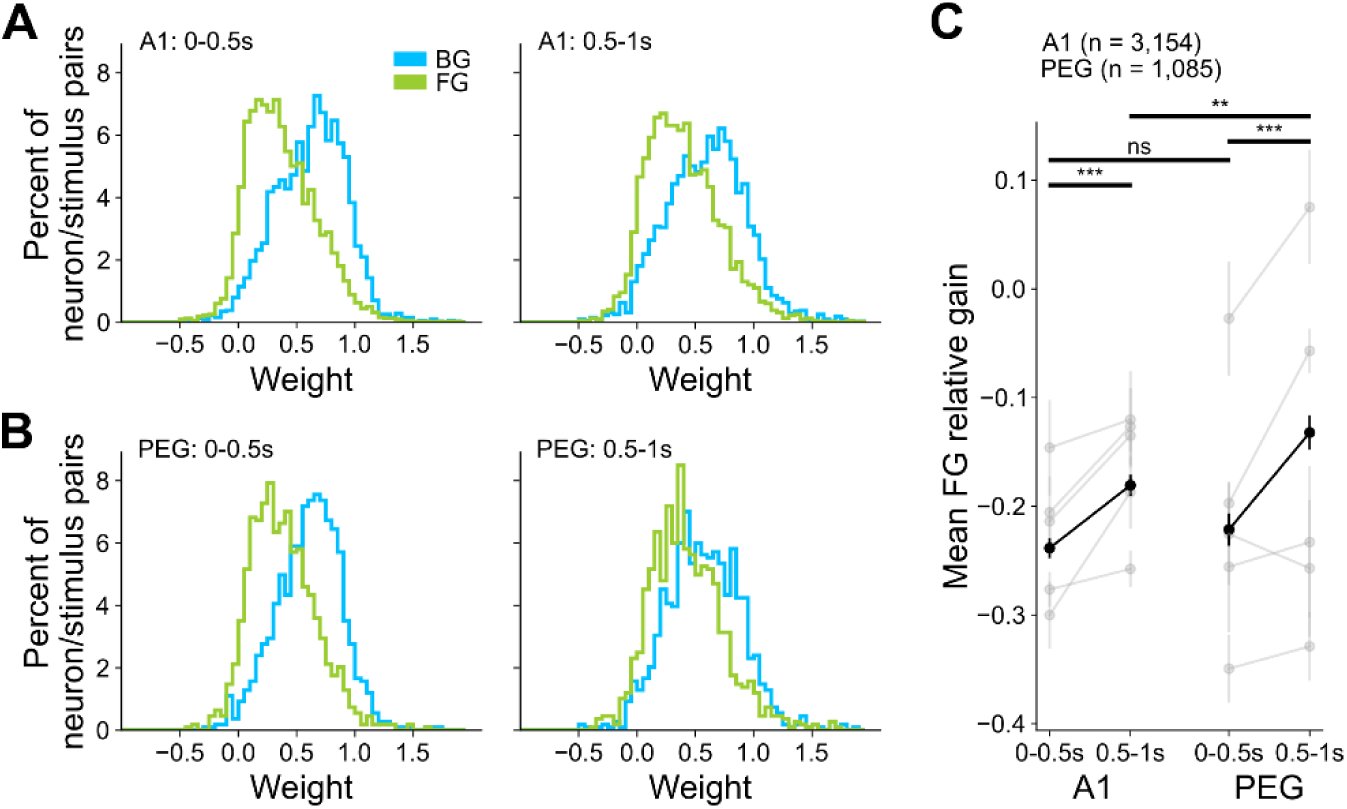
FG suppression decreases after adaptation to a steady state. (A) Histograms compare distributions of BG and FG weights in A1, fit using responses either during 0-0.5s (*left*) or 0.5-1s after onset (*right*). (B) Distributions of BG and FG weights in PEG over the same time windows. (C) Average FG relative gain (RG_FG_) between FG and BG weights, computed for each time window and cortical field. Data for the entire dataset is in black (mean ± S.E.M. across area, Mann-Whitney U rank test, **p < 0.01, ns: not significant; across time windows, Wilcoxon signed-rank test, ***p < 0.001). Data from individual animals is in gray (n = 5).

Thus, both areas show similar FG suppression following sound onset. After a period of adaptation, the amount of suppression in PEG, which lies later in the auditory processing hierarchy, is reduced relative to A1.

### Distinct spectral and temporal sound statistics contribute to FG suppression

Having established a categorical difference in response weights, we next sought to identify distinguishing statistical properties of BGs and FGs that might give rise to FG-specific suppression. While the distinction between BGs and FGs can be intuitive—BG sounds typically contain less behaviorally relevant information^39^—they can also be distinguished by their statistical properties^4,34–37^. We evaluated the extent to which these distinguishing features can explain FG suppression.

For each sound, we measured three spectro-temporal properties previously reported to distinguish BGs and FGs (detailed in *Methods*). Spectral correlation describes how closely power in different spectral bands co-varies^28,47,48^. Noisier sounds are typically less correlated due to the random nature of noise. As such, FG stimuli typically had greater spectral correlation than BG stimuli (BG: 0.177 ± 0.003, FG: 0.497 ± 0.024, p = 6.56e-5, Figure 5A). Temporal variance describes how transient or dynamic a sound is by computing variance in power across each frequency band over time and averaging across frequency^14^. FG sounds tend to contain more transients^35^ and thus have greater temporal variance than BGs (BG: 21.34 ± 5.86, FG: 58.53 ± 14.42, p = 9.80e-7, Figure 5C). Finally, bandwidth describes the frequency range over which a majority of a sound’s power resides. FG sounds tend to have narrower bandwidth than BG sounds (BG: 3.31 ± 0.27, FG: 2.31 ± 0.18, p = 3.00e-3, Figure 5E).

**Figure 5.**
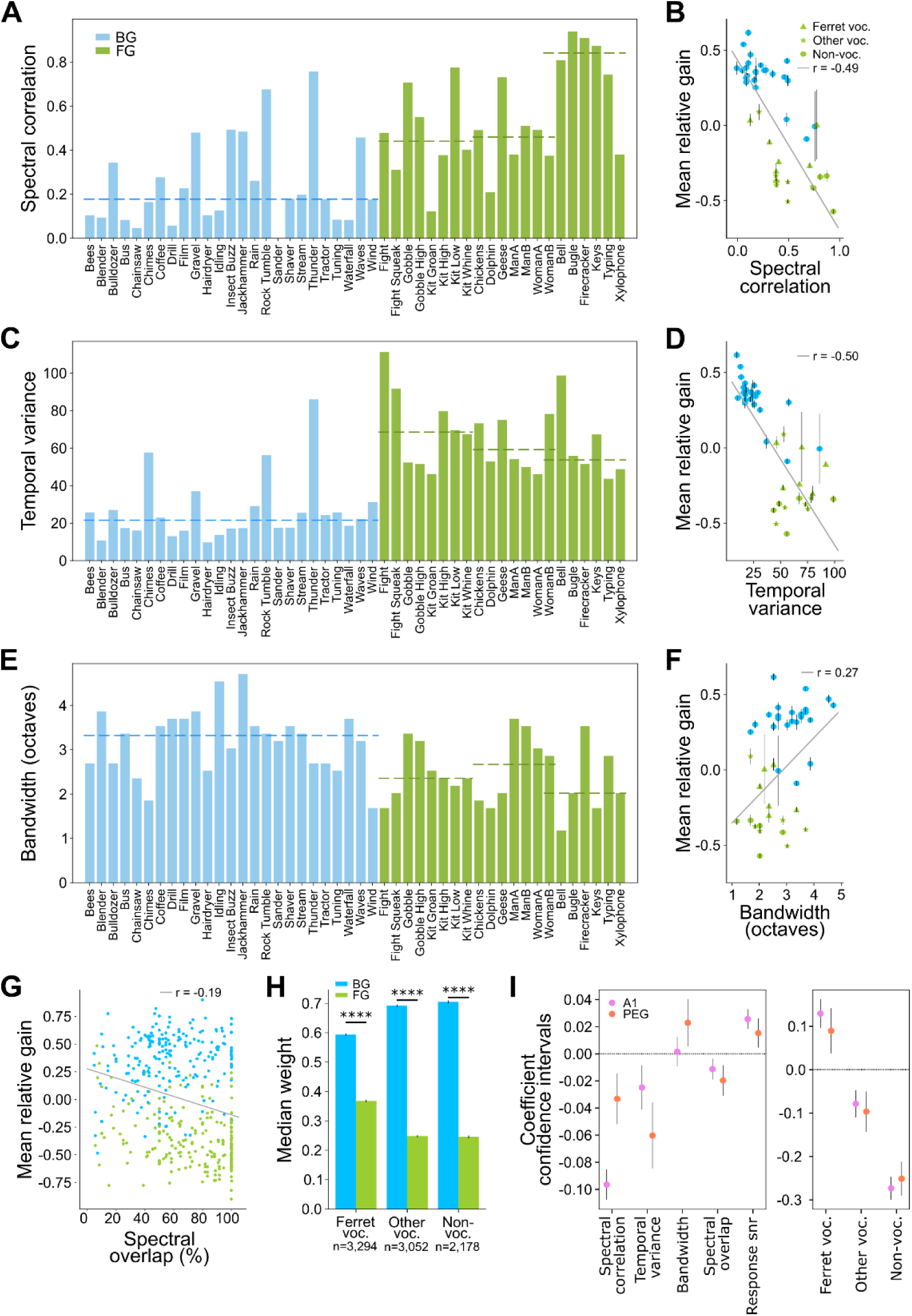
Distinct spectral and temporal sound statistics contribute to FG suppression. (A) Spectral correlation measured for each BG (blue) and FG sound (green). Horizontal dashed lines indicate mean across BGs and subsets of FGs. (B) Scatter plot compares spectral correlation and relative gain for each natural sound in A1 (mean ± S.E.M. per sound; linear regression, r = −0.49, p < 10^-9^). Symbols indicate FG sub-category. (C) Temporal variance of each BG and FG sound. (D) Scatter plot compares temporal variance versus relative gain (r = −0.50, p < 10^-9^), plotted as in (B). (E) Bandwidth of each BG and FG sound. (F) Scatter plot of bandwidth versus relative gain (r = 0.27, p < 10^-9^), plotted as in (B). (G) Scatter plot compares spectral overlap and relative gain for each BG/FG pairing (linear regression, r = −0.19, p < 10^-9^). (H) Average BG and FG weights, after FGs are grouped into vocalization sub-categories (median ± jack-knifed S.E. across neuron/sound pairs, Wilcoxon signed-rank test, ****p < 10^-9^). (I) Relative weight of each sound statistic’s contribution to FG suppression, measured by multivariate linear regression. FG category was treated as a categorical variable; all others were continuous. Response SNR for individual FG and BG sound was included as an input to control for influence of selectivity for the component sounds. The variance of each continuous input was normalized to permit direct comparison of weights. Regression was performed independently for A1 and PEG.

While these statistical properties differ between BGs and FGs on average, their values can vary widely across individual sounds within each category. We next compared how the magnitude of each property affects relative response gain during concurrent sound presentation. In the analysis above, we used FG relative gain (RG_FG_) to describe the relative contribution of FG versus BG to the BG+FG response. We define a complementary statistic, BG relative gain (RG_BG_), to describe the relative contribution of BG to the BG+FG response:

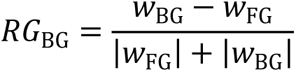

We can now describe the relative gain of any sound, BG or FG, using a single relative gain (RG) metric. As such, RG > 0 indicates that the referenced BG or FG sound is relatively enhanced by a paired sound, and RG < 0 indicates that the referenced sound is relatively suppressed by a paired sound. We measured the relationship between each sound statistic and RG, averaged across neurons and paired stimuli. In A1, spectral correlation was negatively correlated with RG (r = −0.49, p < 10^-9^, Figure 5B), temporal variance was negatively correlated with RG (r = −0.50, p < 10^-9^, Figure 5D), and bandwidth was positively correlated with RG (r = 0.27, p < 10^-9^, Figure 5F). Thus, differences in RG can be attributed to quantitative sound properties rather than the broad BG versus FG categorization. Similar relationships were observed in PEG (Figure S4A).

The properties described above characterize each sound in isolation. We also considered the possibility that the degree of overlap could explain the interaction between sounds. Spectral overlap, or the extent to which the spectral bandwidths of two sounds are matched, has been shown to affect the perception and encoding of concurrent sound sources^49,50^. To calculate spectral overlap, we identified the range of frequencies used to measure bandwidth for each sound. Overlap of sound A with sound B is then defined as the percent of sound A’s bandwidth that overlapped with sound B’s bandwidth (note that this value is not commutative). In both A1 and PEG, there was a small negative correlation of spectral overlap with RG (A1: r = −0.19, p < 10^-9^, Figure 5G; PEG: r = −0.23, p < 10^-9^, Figure S4B).

In addition to differences in spectro-temporal properties distinct between BG and FG sounds, we hypothesized that FG sub-categories contain inherent ethological salience to ferrets and thus may be represented differently. We divided FGs into three categories that might have different significance: 1. ferret vocalizations, 2. vocalizations by other species, and 3. non-vocalizations (Figure 5A, *green bars*). While all three of these categories were significantly suppressed (p < 10^-9^), ferret vocalizations were the least suppressed (Figure 5H, Figure S4C), possibly reflecting their inherent behavioral relevance.

These observations together suggest that a combination of statistical properties can explain FG suppression. At the same time, these properties can be correlated, as is the case of spectral bandwidth and spectral overlap. Thus, to isolate their contributions we performed a multivariate regression to predict RG based on a combination of sound statistics—spectral correlation, temporal variance, bandwidth, spectral overlap, FG category—and baseline response amplitude. Regression coefficients largely reflected analyses of individual statistics described above (Figure 5I). Similarly, effects of FG category also persisted, indicating that their differences could not be explained by sound statistics alone. Interestingly, while the effect of all tested factors was largely equal in magnitude between areas, in A1 the effect of higher spectral correlation predicting lower RG was substantially greater than that of PEG. Conversely, temporal variance showed a stronger predictive effect of lower RG in PEG than A1, but with lower magnitude. The broad effect of sound statistics on RG and the unique effects of spectral and temporal statistics between areas prompted further, direct investigation of these sound statistics.

### Degraded stimuli confirm the role of spectro-temporal properties in FG suppression

To directly test the role of spectro-temporal sound statistics on relative suppression of FG responses, we generated synthetic BG and FG sounds in which temporal and/or spectral modulation features were selectively degraded while preserving other features^32^. For a subset of recordings, synthetic stimuli were generated to match each natural BG/FG pair and presented on randomly interleaved trials. All stimuli were matched in their power spectrum (i.e., “cochlear”- level statistics). Four synthetic conditions were tested: 1. Preserved spectral and temporal statistics (spectro-temporal: ST), 2. Preserved spectral and degraded temporal statistics (spectral: S), 3. Preserved temporal and degraded spectral statistics (temporal: T), and 4. Degraded spectral and temporal statistics (cochlear: C, Figure 6A). Pairs were always presented from the same natural/synthetic category.

**Figure 6.**
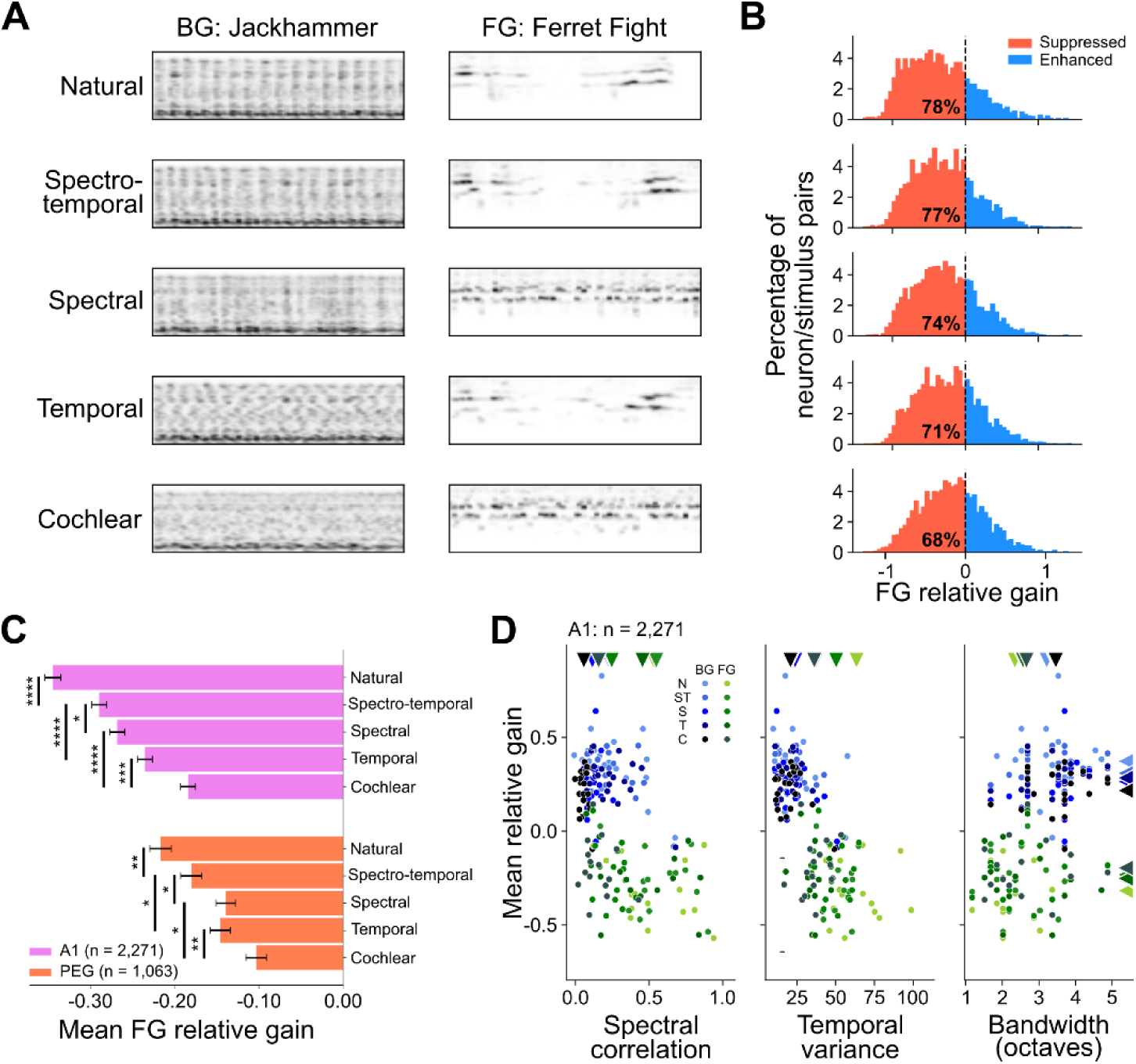
Degradation of natural spectro-temporal properties successively decreases FG suppression in A1. (A) Spectrograms of BG and FG natural sounds and the progression of four synthetic conditions. Labels at left indicate the statistical properties (spectral, temporal modulations) preserved in each example. The cochlear condition lacks natural spectral and temporal modulation statistics. (B) Histograms of FG relative gain (RG_FG_) for each synthetic condition across the matched subset of A1 neurons presented synthetic stimuli (n = 2,271). Red bars and percentages indicate neuron/stimulus pairs showing FG suppression (RG_FG_ < 0). (C) Average RG_FG_ for each synthetic condition in A1 and PEG (mean ± S.E.M. across synthetic conditions, Wilcoxon signed-rank test, *p < 0.05, **p < 0.01, ***p < 0.001, ****p < 10^-9^). (D) Scatter plots compare sound statistics and RG for all natural and synthetic conditions in A1. Saturation indicates level of degradation (N: natural, ST: spectro-temporal, S: spectral, T: temporal, C: cochlear). Triangles along margins indicate average statistic value and RG for each synthetic group.

The relative suppression of FG responses (measured by FG relative gain) decreased successively with the removal of natural spectral and temporal modulations (Figure 6C). There was a significant decrease in FG suppression between the natural sounds and the spectro-temporal synthetic, indicating that higher order statistics and relationships present in natural sounds also contribute to the suppression of FG sounds. Moreover, even in the most degraded, cochlear synthetic condition, FG suppression was not eliminated, confirming that bandwidth differences or simpler “cochlear”-level statistics also contribute to the effect. Both A1 and PEG showed similar successive decreases in FG suppression, but with less overall suppression in PEG for each condition. There were also small differences between areas. Consistent with regression coefficients of individual statistics (Figure 5I), degradation of spectral statistics in A1 had a much greater effect decreasing FG suppression than did the degradation of temporal statistics. Likewise, the degradation of spectral statistics in A1 decreased FG suppression more than in PEG, where degradation of spectral or temporal statistics produced an equal effect (Figure 6C).

Further, after removing specific stimulus identity, incremental degradation of sound statistics caused the relative response to BG and FG to converge in A1 (Figure 6D). These results confirm that multiple statistical properties confer unique attributes to FG sounds that lead to their preferential suppression in AC.

### Spatial and level relationships of sounds influence FG-specific suppression

In all results described thus far, stimuli were presented from a single location 30° contralateral to the recorded brain hemisphere *(contra*BG*/contra*FG). To explore the effect of spatial location on FG-specific suppression, we varied the location of each stimulus between the contralateral position and a second location 30° ipsilateral to the recorded brain hemisphere. This defined three additional spatial configurations: *ipsi*BG/*contra*FG, *contra*BG/*ipsi*FG, and *ipsi*BG/*ipsi*FG. Given the tendency of AC to preferentially encode contralateral stimuli^51,52^, we expected decreased FG suppression when the BG was presented ipsilaterally. Conversely, we expected increased FG suppression when the FG was presented ipsilaterally. Indeed, A1 and PEG both showed significant decreases in FG suppression in the *ipsi*BG/*contra*FG condition (A1: p < 10^-9^, PEG: p < 10^-9^) and significant increases in FG suppression in the *contra*BG/*ipsi*FG condition (A1: p <10^-9^, PEG: p <10^-9^, Figure 7A). The *ipsi*BG/*ipsi*FG condition showed a modest decrease in FG suppression in A1 (p = 0.039) and no significant difference in PEG (p = 0.891).

**Figure 7.**
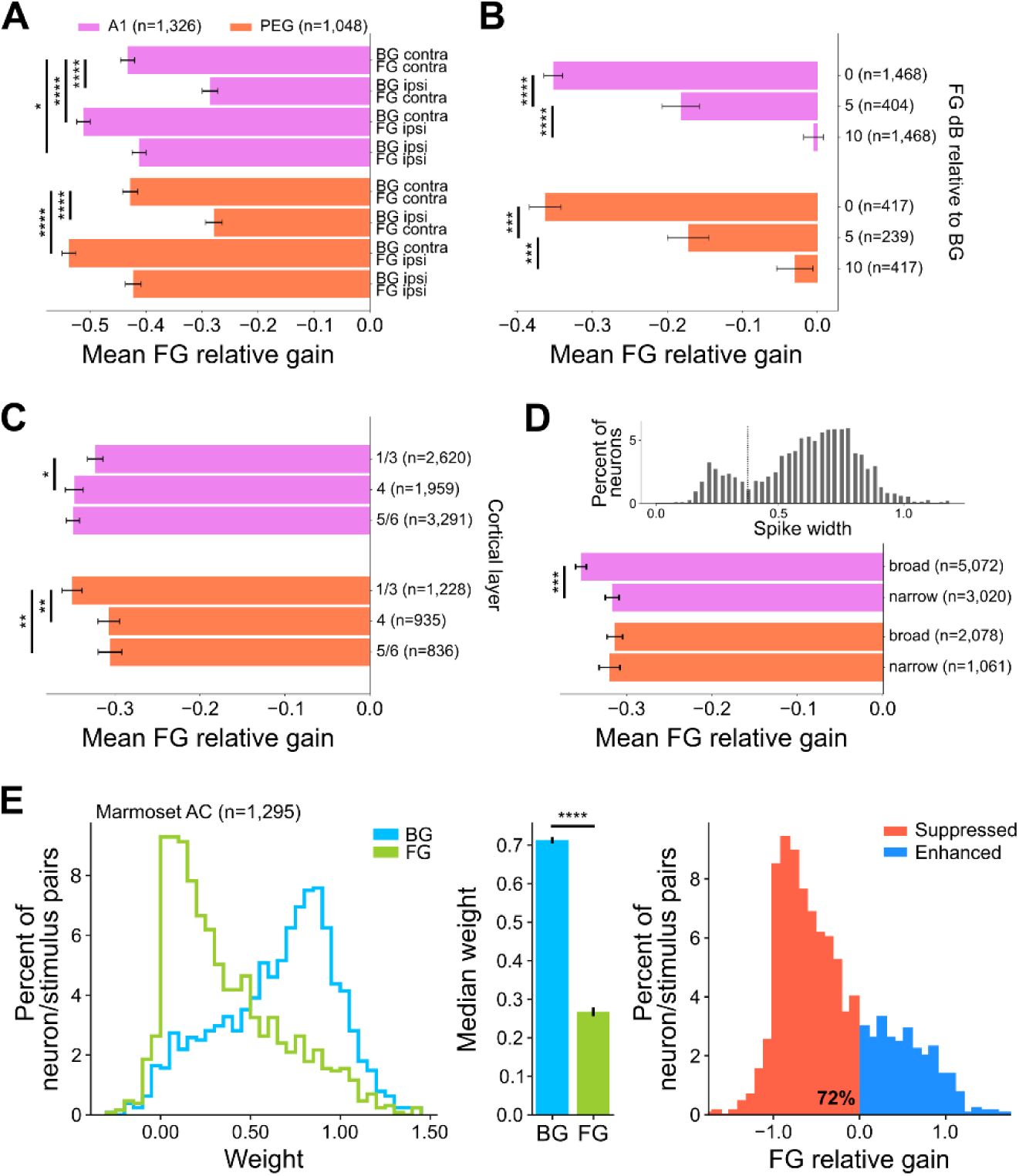
FG suppression across multiple experimental conditions. (A) Average FG relative gain (RG_FG_) for combinations of contralateral (contra) and ipsilateral (ipsi) BG and FG presentations, grouped by cortical field. (A1: n = 1,326, PEG: n = 1,048, mean ± S.E.M, Wilcoxon signed-rank test, *p < 0.05, ****p < 10^-9^). (B) Average RG_FG_ for variable FG sound level (dB), relative to BG, plotted as in (A) (***p < 0.001). N for each group indicated in figure. (C) Average RG_FG_ for neurons grouped by cortical layers, plotted as in (A) (Mann-Whitney U rank test, *p < 0.05, **p < 0.01). (D) (*upper*) Histogram of spike width in A1 shows bimodal peaks corresponding to putative inhibitory (narrow width) and excitatory (broad width). (*lower*) Average RG in for neurons grouped by spike width (Mann-Whitney U rank test, ***p < 0.001). (E) (*left*) Histogram of BG and FG weights for all neuron/sound pair combinations recorded in marmoset AC. (*middle*) Average FG and BG weights (median ± jack-knifed S.E. across neuron/sound pairs, n=1,295, Wilcoxon signed-rank test, ****p < 10^-9^). (*right*) Histogram of RG_FG_ in marmoset AC, where negative RG_FG_ values (red) indicate neuron/sound pairs that show FG-specific suppression (72%).

Varying sound location affected the relative loudness of sounds reaching each ear but also engaged differential circuits for spatial coding. To directly test the effect of relative sound level on responses to the concurrent stimuli, we returned to the *contra*BG*/contra*FG configuration and instead varied the level of FG relative to BG successively from the original 0dB SNR to 5dB and 10dB. Increasing SNR reduced FG suppression (Figure 7B). In the 10 dB SNR condition, RG_FG_ was not statistically different from 0 in A1 or PEG (A1: p = 0.695, PEG: p = 0.214). Thus, while FG suppression only weakly depends on a neuron’s responsiveness to individual BG and FG sounds (Figure 5I), the degree of suppression does depend on overall input strength of each stimulus into the recorded brain area.

### Limited influence of cortical depth and cell type on FG-specific suppression

Our electrode arrays were of sufficient length to acquire units across multiple cortical depths. We used current source density analysis (see *Methods*) to classify units according to their cortical layer (layer 1/3, supragranular; layer 4, granular; layer 5/6 infragranular). In general, differences across layers were small. In A1, there was a slight increase (p = 0.036) in FG suppression between layer 1/3 and layer 4. In PEG, FG suppression in layer 4 and 5/6 were modestly decreased relative to layer 1/3 (layer 4: p = 5.82e-3, layer 5/6: p = 9.23e-3, Figure 7C).

In addition, we classified units as broad- or narrow-spiking based on the width of the spike waveform. We found a characteristically bimodal distribution of spike widths (Figure 7D, *upper*), with a majority of units in the broad-spiking category (A1: 734/1,068, 68.7%, PEG: 342/470, 72.8%). In A1, but not PEG, FG suppression was reduced for narrow spiking compared to broad-spiking units (A1: p = 3.60e-5, PEG: p = 0.784, Figure 7D, *lower*).

### FG-specific suppression also occurs in primate AC

Ferrets are evolutionarily distant from humans, and species differences could explain why FG suppression has not been reported for humans. To determine if the unexpected FG suppression was observed in an animal more closely related to humans, we performed the basic BG/FG contrast while recording single unit activity in AC of the common marmoset (*Callithrix jacchus*). Marmosets are a new world monkey with extensive vocal communication behaviors and AC similar in structure to humans^53–57^. Data from two marmosets recapitulated our observation of strong FG suppression in AC (*w_BG_:* median = 0.714 ± 0.008, *w_FG_:* median = 0.268 ± 0.012, p < 10^-9^, n = 1,295, Figure 7E).

## Discussion

We examined single-unit representations of concurrent natural background (BG) and foreground (FG) sounds in ferret auditory cortex (AC). Presenting BG textures concurrently with dynamic FG stimuli resulted in unexpected FG-specific suppression in both primary (A1) and secondary AC (PEG). This suppression was dependent on statistical features unique to natural FG sounds, and the degradation of natural statistics resulted in a reduced level of suppression. The degree of FG suppression was also reduced following prolonged stimulation, and it varied with manipulation of spatial location and signal-to-noise levels. Across all conditions tested, however, the majority of neurons showed suppression of the FG response relative to the concurrent BG response. A similar suppression was also observed in marmoset AC, indicating that the phenomenon occurs across species. An enhanced representation of natural background stimuli at early stages of auditory processing may support grouping of sound features into perceptual objects, enabling the subtraction of complex noise and producing enhanced representation of behaviorally relevant FG sounds at later stages of processing^8,14,15^.

### Representation of natural foreground and background stimuli in auditory cortex

Previous studies measuring local field potential (LFP) and BOLD (fMRI) in human superior temporal gyrus (STG) have reported noise-invariant representations of behaviorally relevant foreground sounds in the presence of synthetic^8,15^ and natural^14^ background noise. Data from primary AC (A1) has been more limited. One functional imagining study of natural background/foreground contrasts found that noise-invariance may not be present at this earlier processing stage, instead suggesting noise-robustness emerges as information travels from A1 to non-primary AC^45^. At the same time, evidence for noise invariance has been shown as early as A1 using stimulus reconstruction methods for recordings in animal models^13,38,39^. This work has focused on foreground coding in the presence of static, synthetic background sounds, which may explain discrepancies with the current study. As our data demonstrate, foreground responses are less suppressed by synthetic backgrounds that lack natural spectral and temporal statistics (Figure 6C). It is also important to note that reconstruction methods may produce a robust representation of the foreground in the population activity, but they do not provide direct measures of the information is preserved about background noise^40–42^. Previous work that did use a natural background/foreground contrast to measure evoked sound responses has found that natural background stimuli are sometimes represented in A1^43^, a phenomenon that we have shown to be the dominant pattern in AC.

Previous work has left open questions about how noise invariance emerges in the auditory processing hierarchy. Our study directly addressed this uncertainty by pairing diverse natural background and foreground sounds to examine what properties of these statistically complex sounds might give rise to invariance at the single-unit level. Surprisingly, our results showed widespread foreground suppression in single-unit representations in both A1 and secondary AC. Prior work with static background noise has suggested that foreground invariance may emerge by sculpting away inputs in spectro-temporal channels that fall out of band from the FG signal^13,39^. However, the spectro-temporal features of natural BG sounds can overlap with those of the FG. Theories of grouping these complex sounds suggest that their component features must be identified based on coherent activity across subpopulations of neurons tuning to features from to one or the other sound source^21,36^. An overrepresentation of the BG in A1 may support grouping of these features so that they can be subtracted to produce an enhanced FG representation in downstream areas.

### Natural sound statistics impact background/foreground contrasts

Our results emphasize the role of the statistical properties of natural sounds in shaping representations of concurrent backgrounds and foregrounds. Background and foreground stimuli could be readily distinguished by multiple spectral and temporal properties. The magnitude of these statistics predicted the degree of foreground suppression for individual background/foreground pairs (Figure 5). Some of these effects were consistent with expectations from prior work. For example, background sounds tend to have broader bandwidth, which can provide energetic masking of the narrowband foregrounds^49^. Consistent with this idea, our results showed background sounds with broader bandwidth tended to produce greater foreground suppression. However, other relationships were unexpected. For example, foreground sounds tend to have high temporal variance, consistent with their transient dynamics. Transient sounds are typically expected to pop out perceptually form continuous noise^13–15,38^; however, foreground sounds with greater temporal variance were actually more suppressed in A1 and PEG.

Confirming the influence of sounds statistics on foreground suppression, our results show that incremental degradation of that natural features successively decreased the degree of suppression (Figure 6C). Importantly, suppression was even reduced between fully natural sounds and synthetic sounds that preserved all the measured spectral and temporal properties. Thus, fully natural sounds contain additional, unmodeled properties that further engage foreground suppression.

The mechanisms weighing the relative response to foreground versus background may depend on experience and behavior. For the subset of foreground sounds that were ferret vocalizations, we observed less suppression than other foregrounds, a difference that could not be explained by any of the measured sound statistics. Conspecific vocalizations are likely to have strong behavioral salience, and prior studies of streaming have described top-down contributions to streaming whereby behaviorally relevant stimuli are selectively enhanced^8–11,13^. Alternatively, there may be additional, unmodeled statistical properties of vocalizations that support their relatively enhanced representation in A1. Experiments in behaving animals can determine if behavior salience impacts the level of foreground suppression.

### Mechanisms driving the neural representation of concurrent sounds

Our analysis of the interaction between stimulus statistics and relative gain provides insight into possible circuit mechanisms that shape responses to concurrent sounds. Even the most degenerate synthetic stimuli, which lack natural spectral and temporal modulation, still produce suppressed foreground responses. These synthetic background sounds maintained a broader, more uniform power spectrum than foregrounds, which is likely to activate a wide range of tonotopic channels. Lateral inhibition is a widespread property of cortex^58–61^, and broadband stimuli are likely to engage widespread inhibition that could suppress the foreground response. This hypothesis is bolstered by our results showing that foregrounds are suppressed even in neuron/sound pairs where the background did not elicit a significant response (Figure S3B), suggesting that background stimuli evoke subthreshold inhibition that acts on responses to the concurrent foreground. More generally, the weak relationship between responses to the isolated sounds and relative gain suggests that foreground suppression is driven by network-level activity evoked by the background (Figure S3A).

Our experiments investigating the dynamics of adaptation to concurrent sounds provides further evidence for the importance of network activity in shaping responses. Studies of speech coding in noise have shown that human AC responses adapt to the introduction of a noisy background over a few hundred milliseconds^14,62^, a timescale broadly consistent with the single-unit data reported here. Adaptation time following the onset of an interrupting background sound substantially exceeded that of an interrupting foreground (Figure 3C, 3E). Increased adaptation times to background sounds is likely a consequence the background activating a larger portion of the network and therefore requiring longer to reach a steady state, compared to a narrowband foreground.

Analysis of response dynamics also revealed differences across the processing hierarchy. Both A1 and PEG showed similar levels of foreground suppression immediately after sound onset, but PEG showed less suppression in the latter half of the 1-sec stimulus (Figure 4). This release from suppression may reflect a step toward enhanced foreground representation at later stages. Suppression in A1 versus PEG also differed in its dependence on sound statistics. Sounds with large spectral correlation were strongly suppressed in A1 while spectral correlation had relatively little effect in PEG (Figure 5I). Previous work has shown that neurons in secondary AC fields are more selective for complex spectral patterns than A1^63–66^. A neuron tuned to the complex spectral pattern that defines a foreground stimulus may be less susceptible to suppression than an A1 neuron tuned to only a component of its spectrum.

In summary, the representation of concurrent background and foreground sounds in early AC selectively suppresses responses to foreground sounds. This suppression is not obviously consistent with enhanced foreground representations reported at later stages of auditory processing. Instead, these results highlight the open question of how mixed representations in the periphery are grouped and segregated in order to produce an enhanced foreground percept. Our findings suggest a new model for this computation. A strongly enhanced representation of a background sound in A1 may support grouping of the correlated features that comprise that sound^21,36^. The correlated activity of neurons encoding these features can be grouped and subtracted in downstream areas. By removing temporally coherent background features, the resulting activity can represent component features of the foreground, even those that overlap with the background at different times in the stimulus.

## Material and Methods

### Surgical Procedures

All procedures were approved by and performed in accordance with the Oregon Health & Science University Institutional Animal Care and Use Committee (IACUC) and conform to the standards of the Association for Assessment and Accreditation of Laboratory Animal Care (AAALAC) and the United States Department of Agriculture (USDA). Five neutered, descented young adult male ferrets were obtained from a supplier (Marshal Farms). In each animal, sterile head-post implantation surgeries were performed under anesthesia to expose the skull over the auditory cortex (AC) and permit head-fixation during neurophysiology recordings. Surgeries were performed as previously described^67–69^. In brief, two stainless steel head posts were anchored along the midline using light-cured bone cement (Charisma, Kulzer). To improve implant stability, 8-10 stainless self-tapping set screws were mounted in the skull. Layers of bone cement were used to build the implant to a final shape amenable to neurophysiology and wound margin care, which included frequent cleaning and sterile bandaging. Following a two-week recovery period, animals were habituated to head-fixation and auditory stimulation.

Additional data was collected from AC of two spayed and neutered adult marmosets (one female, one male) obtained from the University of Utah. All surgical and experimental procedures performed on marmosets were the same as for ferrets.

### Acoustic Stimuli

Digital acoustic signals were transformed to analog (National Instruments), amplified (Crown), and delivered through a free-field speaker (Manger) placed 80 cm from the animal’s head, 0 ° elevation, and 30 ° contralateral to the recording hemisphere. Stimulation was controlled using custom MATLAB software (https://bitbucket.org/lbhb/baphy) and all experiments took place inside a custom double-walled sound-isolating chamber (Professional Model, Gretch-Ken).

Auditory stimuli consisted of a pool of 70 natural sound excerpts, each 1 s in length, curated and segmented to contain power immediately after onset. Sounds were divided into two ethological categories, backgrounds (BGs) and foregrounds (FGs), based on simple statistics that, respectively, produce the percept of a sound texture or dynamic transient. Sounds were root mean square (RMS) normalized to impose a 0 dB signal-to-noise ratio (SNR) between BG and FG categories for most experiments. Sound level was calibrated so that individual sounds were presented at 65 dB SPL. All excerpts were presented in isolation to each recording site, and the 3-5 sounds from each category that evoked the largest average multi-unit response were selected for experiments. Selected sounds were combinatorically paired across categories to create 9-25 unique BG/FG combinations. The full set of isolated and concurrent sounds was presented in random order 10-20 times per recording.

We tested several variations of BG/FG combinations:

#### Dynamic sound onsets

To study dynamics of adaptation to concurrent stimulation (Figure 3), we presented natural pairs in which either FG or BG was truncated to its 0.5-1s half while the paired sound was played in full (BG+hFG or hBG+FG, Figure 3A, 3D). FG sounds used in the dynamic conditions were selected so that they contained energy both at 0 and 0.5 sec.

#### Binaural stimulation

In our default experimental configuration, all sounds were presented from a single speaker 30° contralateral to the recorded brain hemisphere (*contra*BG/*contra*FG). For binaural stimulation, we added a second speaker 30° ipsilateral to the recording hemisphere. In these recordings, BG and FG positions were varied, defining three additional spatial configurations: *ipsi*BG/*contra*FG, *contra*BG/*ipsi*FG, and *ipsi*BG/*ipsi*FG.

#### Variable SNR

In a subset of experiments, we the RMS power FG relative to BG by 5 dB and 10 dB to produce instances where FG had greater SNR than BG.

### Neurophysiology

To prepare for neurophysiological recordings, a small craniotomy (0.5-1mm) was opened over AC. Recording sites were targeted based on tonotopic maps and superficial skull landmarks^64,70^ identified during implantation surgery. Initially, tungsten microelectrodes (FHC, 1-5MΩ) were inserted into the craniotomy to characterize tuning and response latency. Short latency responses and tonotopically organized frequency selectivity across multiple penetrations defined the location of primary auditory cortex (A1)^70^, whereas secondary auditory cortex (posterior ectosylvian gyrus, PEG) was identified as the field ventrolateral to A1. The border between A1 and PEG was identified from the low-frequency reversal of the tonotopic gradient.

Once a cortical map was established, subsequent recordings performed using two different electrode configurations. Experiments in animals 1-3 used 64-channel silicon electrode arrays, which spanned 1.05mm of cortical depth^71^. Experiments in animals 4-5 recorded from 960-channel Neuropixels probes^72^. Typically, about 150 of the 384 active channels spanned the depth of AC, as determined by current source density analysis (see below). Data were amplified (RHD 128-channel headstage, Intan Technologies; Neuropixels headstage, IMEC), digitized at 30 KHz (Open Ephys)^73^, and saved to disk for further analysis. Spikes were sorted offline using Kilosort2 (https://github.com/MouseLand/Kilosort2)^74^, with spike sorting results manually curated in phy (https://github.com/cortex-lab/phy). A contamination percentage was computed by measuring the cluster isolation for each sorted and curated spike cluster, which was classified as a single unit if contamination percentage was less than or equal to 5%. Clusters above 5% were classified as multi-unit and excluded from analysis. Neurophysiology recordings were performed in animals in a passive state while head fixed and unanesthetized, with sessions typically lasting 4-6h.

### Inclusion Criteria

Evoked activity was measured using the peri-stimulus time histogram (PSTH) response, averaged across 10-20 sound repetitions and sampled at 100Hz. To ensure only sound-responsive units were included in analyses, we calculated a signal-to-noise ratio for each stimulus/neuron pair based on the ratio of the PSTH response to the standard deviation of the response across repetitions^75^. Stimulus-neuron pairs with SNR ≥ 0.12 were considered sound-responsive and included for analysis (Figure S2A, S2D). Neuron/stimulus pairs where responses to both BG and FG in isolation exceeded SNR threshold were categorized as responsive to both sounds (BG^+^/FG^+^- A1: 9,057/29,445, 30.8%, PEG: 3,838/14,936, 25.7%) and comprise the dataset used in most analyses. Other stimulus/neuron pairs were categorized as responsive to only one sound (BG^0^/FG^+^ - A1: 3,117/29,445, 10.6%, PEG: 1,498/14,936, 10.0%; BG^+^/FG^0^ - A1: 2,186/29,445, 7.4%, PEG: 954/14,936, 6.4%) or unresponsive (BG^0^/FG^0^ - A1: 15,085/29,445, 51.2%, 8,646/14,936, PEG: 57.9%). Unresponsive neuron/sound pairs were excluded from all analyses.

Within responsive neuron/sound pairs, outlier instances for which BG or FG weights were less than −0.5 and greater than 2 were also excluded (A1: 9/9,057, PEG: 10/3,838). Further, only instances where the linear model fit well (r ≥ 0.4, A1: 8,524/9,048, 94.2%, PEG: 3,492/3,828, 91.2%) were included. Among the minority of neuron/sound pairs excluded based on model accuracy, response weights trended with the r ≥ 0.4 data (Figure S2B, S2E).

### Stimulus Statistics

To calculate sound stastistics, natural sound spectrograms were generated from .wav files using 10 ms time bins and 48 frequency channels, log-spaced 0.1-24kHz.

#### Bandwidth

quantified the range of frequencies that contained the majority of power in the spectrogram. A power spectrum was calculated by averaging each spectrogram over time and computing the cumulative sum across frequency. Bandwidth was determined by identifying the frequency range, in octaves, between 15% and 85% of the total.

#### Spectral correlation

was describing how closely power across spectral bands co-varies. The correlation coefficient (Pearson’s R) was computed over time between each pair of frequencies in the calculated bandwidth range, and averaged across frequency pairs.

#### Temporal variance

was quantified by calculating variance over time in each spectral channel in the calculated bandwidth range and averaging across frequencies. This statistics is also referred to as “temporal non-stationariness"^14^. Greater deviations indicate higher temporal variance, or a more dynamic and transient sound.

#### Spectral overlap

was the fraction of a sound’s bandwidth that overlapped the bandwidth of a concurrently presented sound. Note that because individual sounds varied in bandwidth, this metric is not commutive, as the overlap of A with B is not the same as the overlap of B with A.

### Synthetic Sounds

We generated model-matched, synthetic BG and FG sounds using a published Matlab toolbox^32^. Temporal and/or spectral modulation features were selectively degraded to generate four synthetic conditions: 1. Preserved spectral and temporal statistics, 2. Preserved spectral and degraded temporal statistics, 3. Preserved temporal and degraded spectral statistics, and 4. Degraded spectral and temporal statistics. Synthetic BGs and FGs were paired within each synthetic category, and presentations were interleaved with natural BG/FG pairs.

### Laminar Depth Analysis

We used current source density analysis to classify units by cortical layer (1/3, supragranular; 4, granular; 5/6 infragranular). The local field potential (LFP) signal was generated by lowpass filtering either the raw signal from the 64-channel silicon probe or the LFP signal from the Neuropixel probe below 250Hz using a zero-phase shift (filter-filter method) 4^th^ order Butterworth filter, followed by down-sampling to 500Hz. A custom graphical interface was used to mark boundaries between layers, based on features of average sound-evoked LFP traces sorted by electrode depth (https://github.com/LBHB/laminar_tools). Layer-specific features included the pattern of current source density (CSD) sinks and sources evoked by best frequency-centered broadband noise bursts. Patterns were selected to match auditory evoked CSD patterns seen in AC of multiple species^76–79^. Each unit was assigned a layer based on the boundaries above and below the channel where its spike had largest amplitude.

### Spike Width Classification

We classified neurons as narrow- and broad-spiking based on the average width of the waveform. Width was calculated as the time between the depolarization trough and the hyperpolization peak^80^. The distribution of spike width across neurons was bimodal, and the categorization threshold was defined as the minimum between the bimodal peaks. Filtering properties differed between 64-channel probes and Neuropixels, thus categorization threshold was defined as 0.35 ms and 0.375 ms, respectively.

## Acknowledgements

The authors would like to thank Sam Norman-Haignere for guidance on generation of model-matched synthetic stimuli and members of the David Lab for feedback on data analysis.

## Author contributions

G.R.H. and S.V.D. designed experiments, analyzed data, and wrote the manuscript. G.R.H., J.C.W., L.A.S., and M.L.E. collected the data.

