## Supplemental Figures for "Unexpected suppression of neural responses to natural foreground versus background sounds in auditory cortex"

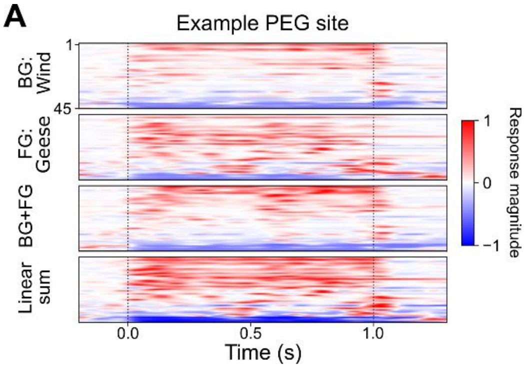


**Figure S1.** Heatmaps show normalized PSTH responses for all units recorded from the same PEG site as the example in Figure 1c (n = 45). Similar to the A1 site in Figure 1d, responses to the BG+FG sound have lower magnitude than the linear sum of responses to the isolated BG and FG sounds for most units.

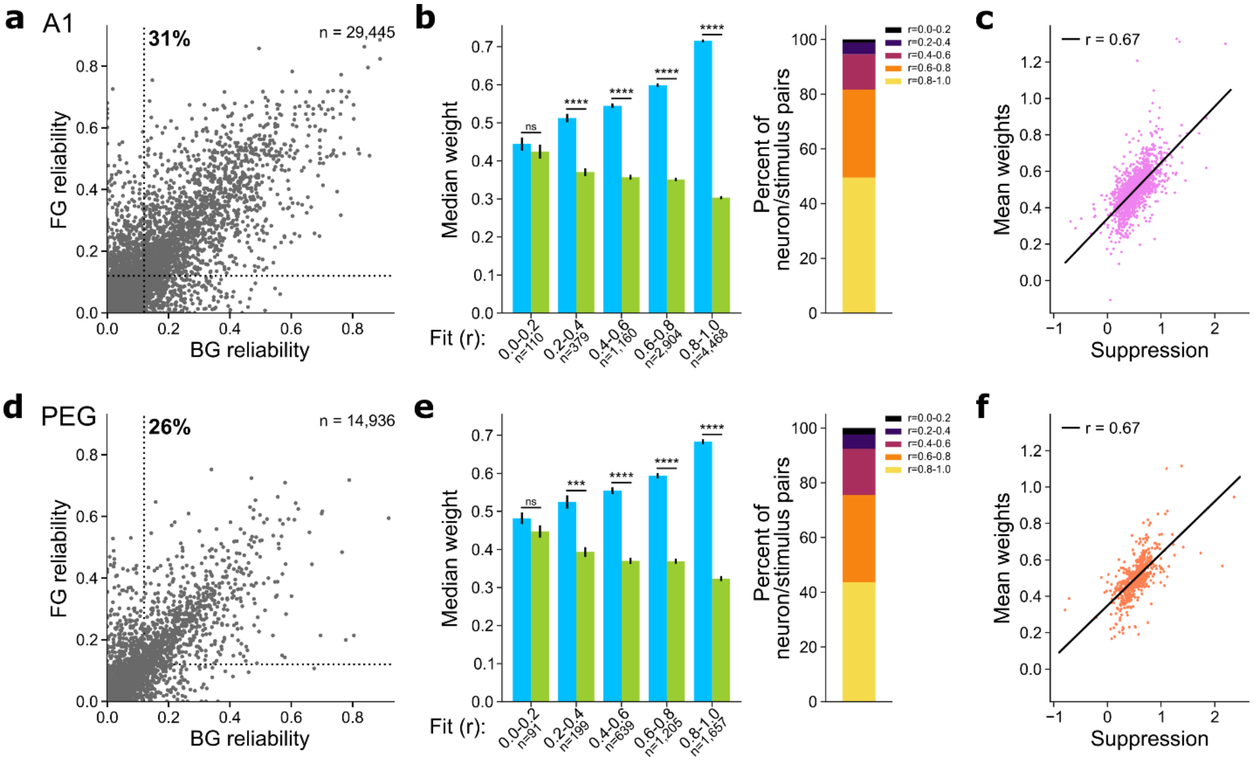


**Figure S2.** Inclusion criteria for neuron/sound pairs. **a** Scatter plot compares reliability (signal power/noise power) of responses to BG and FG stimuli for each neuron/sound pair combination in A1. The threshold for sound responsiveness was set at 0.12 (dotted lines), labeling 31% as responsive to both BG and FG. **b** Median BG and FG weights in A1 grouped by fit accuracy (r) of the weighted linear model (*left*, median ± jack-knifed S.E. across neuron/sound pairs, Wilcoxon signed-rank test, ****$p < 10^{-9}$). (*right*) Percent of total (n = 29,445) neuron/sound pairs in each fit accuracy range, indicating that the majority (94.2%) had r ≥ 0.4, the threshold for inclusion in the subsequent relative gain analysis. **c** Scatter plot compares the direct measure of response suppression versus the mean fitted weight for each neuron/sound pair combination in A1 that met criteria for response reliability and model accuracy (linear regression, r = 0.67, ****$p < 10^{-9}$). **d-f** For PEG data, comparison of (d) response reliability, (e) BG and FG weights grouped by model accuracy (n = 14,936, ***$p < 0.001$, ****$p < 10^{-9}$, r ≥ 0.4, 91.2%), and (f) comparison direct versus model suppression (r = 0.67, ****$p < 10^{-9}$), plotted as in (a)-(c).

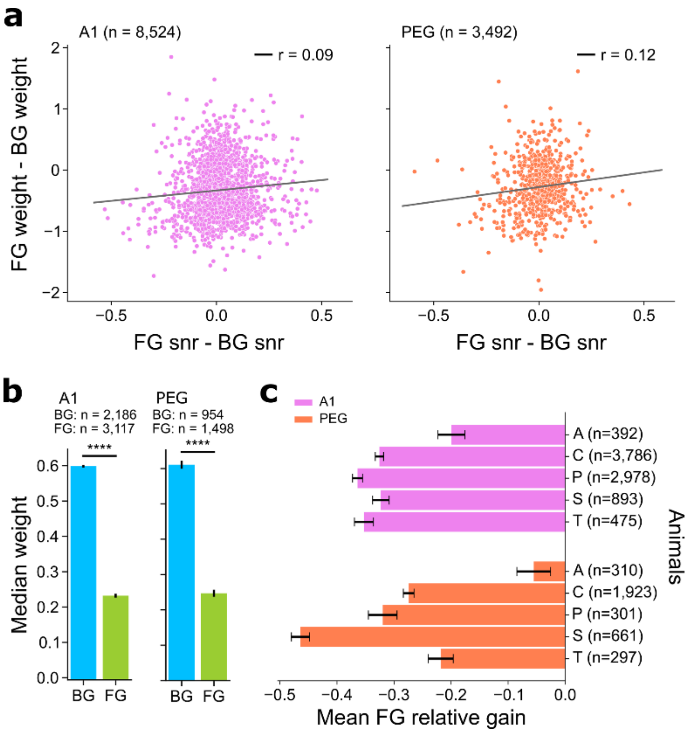


**Figure S3.** FG suppression is reliable and is not simply a function of single-stimulus responsiveness. **a** Scatter plots compare the difference in FG and BG responsiveness versus FG relative gain for each neuron/stimulus pair tested in A1 (*left*, n = 8,524, linear regression, r = 0.09, p = 2.23e-4) and PEG (*right*, n = 3,492, linear regression, r = 0.12, p = 2.00e-3). **b** Average BG and FG weights for neuron/sound pairs where the neuron is only responsive to BG or FG in A1 (*right*, median ± jack-knifed S.E. across neuron/sound pairs, Mann-Whitney U rank test, ****$p < 10^{-9}$) and PEG (*right*, ****$p < 10^{-9}$). Weights are shown for the BG or FG stimulus that evoked a significant response when presented concurrently with a stimulus that did not produce a response, i.e., with zero weight. **c** Breakdown of average FG relative gain ($RG_{FG}$) for each recorded animal (n = 5). $RG_{FG}$ was lower in A1 than in PEG for 4/5 animals.

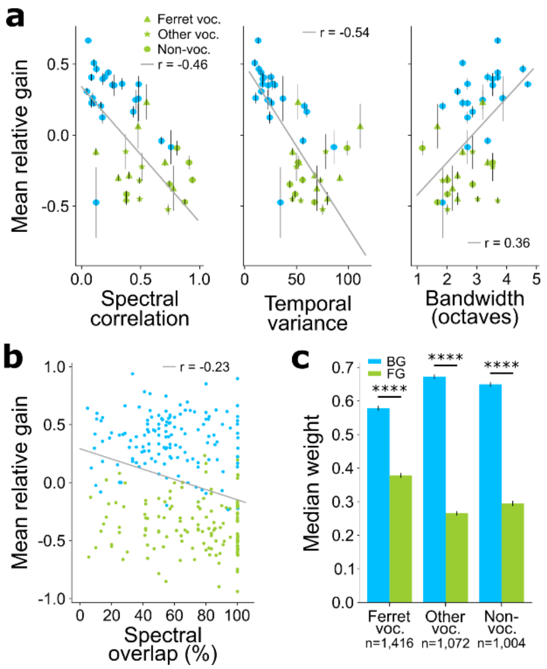


**Figure S4.** Contribution of sound statistics to FG-specific suppression in PEG. **a** Scatter plots compare sound statistics versus relative gain (RG) in PEG, plotted as in Fig. 5 (mean ± s.e.m. across sounds, linear regression). All three statistics show similar patterns to A1: spectral correlation (*left*, $r = -0.46$, $p < 10^{-9}$), temporal variance (*middle*, $r = -0.54$, $p < 10^{-9}$), and bandwidth (*right*, $r = 0.36$, $p < 10^{-9}$). BGs shown in blue; FGs shown in green; symbols indicate FG sub-category. **b** Scatter plot compares spectral overlap and relative gain for each BG/FG pair (linear regression, $r = -0.23$, $p < 10^{-9}$). **c** Average BG and FG weights after grouping FGs into vocalization sub-categories (median ± jack-knifed S.E. across neuron/sound pairs, ****Wilcoxon signed-rank test, $p < 10^{-9}$).
